# Comparative metagenome-assembled genome analysis of *Lachnovaginosum* genomospecies, formerly known as BVAB1

**DOI:** 10.1101/657197

**Authors:** Johanna B. Holm, Michael France, Bing Ma, Courtney K. Robinson, Elias McComb, Rebecca M. Brotman, Jacques Ravel

## Abstract

Bacterial Vaginosis Associated Bacteria 1 (BVAB1) is an uncultured bacterial species found in the human vagina that belongs to the family *Lachnospiraceae* within the order *Clostridiales*, and as its name suggests, is often associated with bacterial vaginosis (BV). In reproductive health, BV is of concern due to its associated risk for HIV, *Chlamydia trachomatis*, and *Neisseria gonorrhoeae* acquisition as well as preterm birth. BVAB1 has been shown to be associated with BV persistence after metronidazole treatment and increased vaginal inflammation, and to confer a higher risk of puerperal infections. To date, no genome of BVAB1 is available, which has made it difficult to understand its disease-associated features. We present here a comparative analysis of seven metagenome-assembled genomes (MAGs) of BVAB1, derived from the cervicovaginal lavages of seven separate women. One metagenome was sequenced with long-read technology on a PacBio Sequel II instrument while the others were sequenced using the Illumina HiSeq platform. The MAGs were 1.5-1.7 Mb long, encoding an average of 1,499 genes and had >98% average nucleotide identity to each other. CheckM-estimated genome completion was 98.2%. We propose to rename BVAB1 to *Lachnovaginosum* genomospecies based on a phylogenetic analysis, and provide genomic evidence that this species may metabolize D-lactate, produce trimethylamine (the chemical responsible for BV-associated odor), and be motile. These MAGs will be a valuable resource and will contribute to our understanding of the heterogenous etiologies of bacterial vaginosis.

## Introduction

Bacterial vaginosis (BV) is a common vaginal infection affecting approximately 21 million or 30% of US reproductive-aged women in 2001-2004, with both African- and Mexican-Americans disproportionately afflicted [2, 3]. The condition is difficult to treat with antibiotics and the rate of recurrence is high [4, 5]. Microbiologically, BV diagnosis is established using Nugent scoring of Gram-stained vaginal smears and is defined by a low abundance of *Lactobacillus* spp. morphotypes, and a high abundance of anaerobic Gram-negative species and/or the presence of clue cells [6]. Clinically, diagnosis of BV is made when 3 of the 4 Amsel’s criteria are met: vaginal pH > 4.5, homogenous vaginal discharge, a positive whiff test, and the presence of clue cells upon wet mount examination [7]. Aside from the burdensome symptoms of vaginal discharge and fishy odor, BV is also associated with increased risk to adverse health outcomes including preterm birth [8], and increased risk of sexually transmitted infections acquisition and transmission, including HIV [9, 10].

A critical step along the path to understanding the ecology and pathogenic potential of a bacterial species is the characterization of its genome. Yet many of these BV associated bacteria have eluded attempts at cultivation, further complicating genome sequencing efforts. BV-associated bacterium 1 (BVAB1) is one such organism for which there is limited genetic information available. Based on 16S ribosomal RNA (rRNA) gene amplicon high-throughput sequencing, BVAB1 belongs to the family *Lachnospiraceae* and is often misidentified as belonging to the genus *Shuttleworthia*. BVAB1 has been associated with both symptomatic and asymptomatic bacterial vaginosis (asymptomatic BV occurs when 3 of 4 Amsel’s criteria are met, but the patient reports no symptoms) [11]. Interestingly, Gram-negative curved rods designated *Mobiluncus* morphotypes on Gram stain in Nugent scoring have been shown to most likely be BVAB1 [12]. Further, the vaginal communities containing this particular bacterial ribotype have been associated with vaginal inflammation and persistent BV [13]. BVAB1 remains uncultured and aside from detection of this species via partial 16S rRNA gene amplicon sequencing, little is known about the metabolism, pathogenic potential, or ecology of BVAB1 in the vaginal environment, especially during BV. Further understanding of the genetic and physiological properties of BV-associated bacteria will help to dissect complex etiology of BV. There currently is no available draft genome for this bacterium.

In this study, we reconstructed seven BVAB1 metagenome-assembled genomes (MAGs) originating from different women with symptomatic and asymptomatic BV. To enable the assembly of high-quality MAGs, we used a sequencing strategy that relied on both Illumina HiSeq and PacBio Sequel II long reads. Based on phylogenetic analysis of full-length 16S rRNA gene sequences obtained from the genomic assemblies, we propose to rename the bacterium *Lachnovaginosum* genomospecies.

## Methods

### Ethics

Samples used in this study were archived cervicovaginal lavages and swabs obtained in accordance with the protocols approved by the University of Maryland Baltimore Institutional Review Board.

### Sample Collection

Vaginal samples used in this study were previously screened and confirmed to contain a high relative abundance (> 70%) of BVAB1 using 16S rRNA gene amplicon sequencing of the V3-V4 regions as previously reported [14]. Cervicovaginal lavages from six participants were collected as part of the NIH Longitudinal Study for Vaginal Flora (LSVF) [15] by washing the vaginal walls with 3 mL sterile, deionized water, and aspiration from the vaginal vault via pipette. Specimen were placed into a plastic tube and frozen at −20°C, and then transferred to −80°C until processing. An additional swab sample collected as part of the UMB-HMP study was used in this study [16]. The swab was self-collected by a participant using a vaginal ESwab and re-suspended into 1mL Amies transport medium (ESwab, Copan Diagnostics Inc.), frozen at −20°C for no more than a week, and transferred to −80°C until analyzed. DNA was extracted from 200 μL of swab or lavage fluid using the MagAttract Microbial DNA Kit (QIAGEN Inc., Germantown MD) automated on a Hamilton Star robotic platform. DNA was eluted in a final volume of 110 μL nuclease-free water.

### Metagenomic library construction and sequencing on the Illumina HiSeq 4000 platform

Metagenomic libraries for the six LSVF samples were prepared using the KAPA HyperPlus Kit (Kapa Biosystems, Wilmington MA) with KAPA Single-Indexed Adapter Kit Set B. A fixed volume (35 μL) of genomic DNA was used as input, and libraries were prepared following the manufacturer’s protocol with modifications based on their amount of input DNA as in **Supplementary File 1**. For samples with 0.5 or 0.2 ng input DNA, the fragmentation enzyme was diluted 1:2 or 1:5 with water. All samples were fragmented at 37°C for 5 min. Adapter concentrations varied according to the input DNA as listed in **Supplementary File 1**, and the adapter ligation was carried out overnight at 4°C for all samples. The post-ligation cleanup was performed with 0.8X Ampure XP beads (Beckman Coulter, Indianapolis IN) and 20 μL of sample was used in library amplification. Amplification library cycles varied by input DNA as listed in **Supplementary File 1**. Post-amplification cleanup was performed with 1x Ampure XP beads; libraries with remaining adapter dimer peaks were cleaned a second time. The final elution was in 25 μL of nuclease-free water. Libraries were run on a Tapestation instrument with a D1000 tape (Agilent, Santa Clara CA) to assess quality and concentration. Each library was sequenced on an Illumina HiSeq 4000 instrument using the 150 bp paired-end protocol.

### Quality filtering, host-read removal, and metagenomic-assembled genome reconstruction

Metagenomic reads were quality filtered using Trimmomatic v0.36 [17] to remove sequencing adapters allowing for 2 mismatches, a palindromic clip threshold of 30, and a simple clip threshold of 10. Bases with quality scores < 3 were removed from the beginning and end of reads, and a 4 bp sliding window which trimmed a read if the average quality score within that window fell below 15. Reads < 75 bp in length were removed. Host reads were detected by mapping to the human genome GRCh38.p12 with Bowtie 2 v2.3.4.1 and default settings [18], followed by removal using samtools v1.9 [19] and bedtools v2.27.1 [20, 21] (see **Supplementary File 2** for specific code). Metagenomic assemblies were produced using MegaHit v1.1.3 [22] with the meta-sensitive setting, and read mapping to assemblies was performed using Bowtie 2 v2.3.4.1 and default settings [18]. Contigs > 2.5 kb were binned based on hierarchical clustering of contig tetranucleotide frequencies and coverage using Anvi’o (version 5.2) [23]. Bins with the highest coverage in the sample which also had the lowest GC content were observed in all samples and were chosen and extracted as candidate BVAB1 MAGs (see **Supplementary File 3** for Anvi’o images of extracted bins). MAGs were annotated using PROKKA v1.13 [24]. MAG completion and contamination estimates were calculated using CheckM v1.0.12 [25]. Phages were detected using PHAST [26].

### Metagenomic library construction and sequencing on the PacBio Sequel II platform

The sample collected from the UMB-HMP study [16] had sufficient DNA content for long-read sequencing using the Pacific Biosciences Sequell II platform. Library pools were prepared with SMRTBell Template Prep Kit 1.0 with barcoded adaptors. Libraries were size-selected on a BluePippen (Sage Science, Beverly, MA) with a cutoff of 5 kb. Sequencing was performed on a Sequel II instrument (Pacific Biosciences, Menlo Park, CA) at the Genomic Resource Center of the University Maryland School of Medicine with a loading at 60 pM.

### Quality filtering, host-read removal, and metagenomic-assembled genome reconstruction

Circular consensus sequencing (CCS) reads were generated using the PacBio CCS application with minPredictedAccuracy=0.99 and the rest of the parameters were set to default, including minimum 3 subread passes. Demultiplexing was done with *lima* (version 1.9.0) using default parameters except for minimum barcode score and min length which were set at 26 and 50 bp respectively. Both tools are part of the SMRTLink 6.0.1 software package with updated CCS version 3.4.1. Human CCS reads were detected and removed using BMTagger [27] and the human genome build 38 patch release 12 (GRCh38.p12). Remaining CCS reads were assembled via Canu v1.8 and the “-pacbio-raw” protocol [28]. Resulting contigs were taxonomically annotated using BLASTN v 2.8.1 [29] and the NCBI non-redundant nucleotide database (updated 2019/05/03) to remove all contigs identified as known vaginal taxa. Contigs were further extended when possible using Circlator 1.5.5 [30]. Based on the GC content of the LSVF MAGs (mean 31%), contigs with median GC content of ≤ 36% were used to produce the UMB-HMP MAG. The contigs were then ordered by decreasing size. The MAG was annotated using PROKKA v1.13 [24]. Phages were detected using PHAST [26]. MAG completion and contamination estimates were calculated using CheckM v1.0.12 [25].

### Metabolomic Data

Metabolomics analysis was conducted as described in [31] by Metabolon Inc. (Morrisville, NC) using 200 μL of lavage sample (sample from LSVF) or 200 μL of a frozen dry swab eluted in 1 ml of PBS (UMB-HMP sample [16]). The abundances of 561 compounds were quantified using Ultrahigh Performance Liquid Chromatography-Tandem Mass Spectroscopy (UPLC-MS/MS). Quantities were corrected for instrument block variability and reported as normalized area-under-the-curve estimates. Figures were generated using ggplot2 [32].

### Comparative genome and phylogenetic analyses

The UMB-HMP MAG served as a reference to order the contigs of the LSVF MAGs using the Mauve Contig Mover [33]. Average nucleotide identities were calculated using FastANI v1.1 with fragment lengths of 1,000 bp which was mean length of coding sequences [34]. A circle plot was constructed with the BLAST Ring Image Generator v0.95 [35] using the PacBio MAG as reference. Maximum-likelihood phylogenetic trees of full-length 16S rRNA gene sequences alignments were generated with ClustalW [36] and FastTree v2.1.5 [37, 38] in Geneious v9.0 (https://www.geneious.com). Trees were rooted with *Fusobacterium nucleatum* (AJ133496) and *Propionigenium modestum* (X54275) [39], and members of the genus *Lachnoclostridium* were included as neighbors [40].

## Results and Discussion

### Participant information

All six participants in the LSVF study were of reproductive age and had high Nugent scores (6-10). Four were diagnosed with asymptomatic Amsel-BV, 1 with symptomatic Amsel-BV, and 1 was not diagnosed with BV [41]. The woman who participated in the UMB-HMP study [16] had a Nugent score of 8 and was diagnosed with asymptomatic BV. The vaginal microbiota of these 7 samples had >60% BVAB1 as defined by 16S rRNA gene V3-V4 amplicon sequencing (**Table 1**) and were used for metagenomic reconstruction of the BVAB1 MAGs.

**Table 1.**
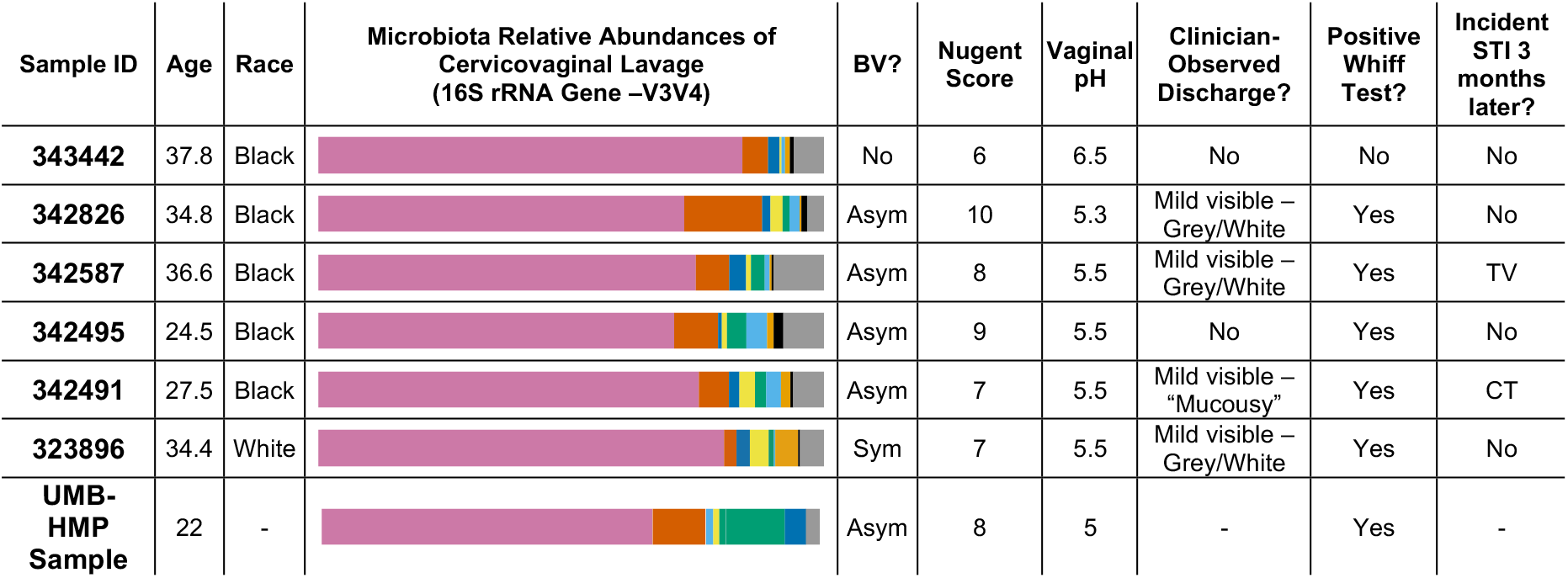
Participants demographics and cervicovaginal lavage microbial compositions for samples used in metagenomic reconstruction of *Lachnovaginosum* genomospecies metagenome-assembled genomes. Microbial compositions indicate from left to right: *Lachnovaginosum* genomospecies (pink), *Gardnerella vaginalis* (darker orange), g_*Megasphaera* (darker blue), *Sneathia sanguinegens* (yellow), g_*Prevotella* (green), *Atopobium vaginae* (lighter blue), *Prevotella amnii* (lighter orange), and Other (gray). Asym: asymptomatic BV; Sym: symptomatic BV.

| Sample ID | Age | Race | Microbiota Relative Abundances of Cervicovaginal Lavage (16S rRNA Gene –V3V4) | BV? | Nugent Score | Vaginal pH | Clinician-Observed Discharge? | Positive Whiff Test? | Incident STI 3 months later? |
| --- | --- | --- | --- | --- | --- | --- | --- | --- | --- |
| 343442 | 37.8 | Black |  | No | 6 | 6.5 | No | No | No |
| 342826 | 34.8 | Black |  | Asym | 10 | 5.3 | Mild visible – Grey/White | Yes | No |
| 342587 | 36.6 | Black |  | Asym | 8 | 5.5 | Mild visible – Grey/White | Yes | TV |
| 342495 | 24.5 | Black |  | Asym | 9 | 5.5 | No | Yes | No |
| 342491 | 27.5 | Black |  | Asym | 7 | 5.5 | Mild visible – “Mucousy” | Yes | CT |
| 323896 | 34.4 | White |  | Sym | 7 | 5.5 | Mild visible – Grey/White | Yes | No |
| UMB-HMP Sample | 22 | - |  | Asym | 8 | 5 | - | Yes | - |

### Overall genomic features

The BVAB 1 draft assembly from the sample sequenced on the PacBio Sequel II comprises of 17 contigs which total 1.57 Mb in size with 31.9% GC content, and is herein termed the “reference MAG”. The BVAB1 MAG constructed using the Illumina HiSeq 4000 metagenomic sequence data were 1.54-1.70 Mb in size (mean: 1.62 Mb) with 27-35 contigs (mean N50: 99,623 bp) and 31.3% GC (**Table 2**). Genome annotation indicated the presence of four complete rRNA operon copies in the reference MAG. Partial rRNA operons were observed the Illumina-based MAGs.

**Table 2.** *Lachnovaginosum* genomospecies genome draft assembly characteristics. Completion and contamination evaluated by CheckM [1].

| Genome ID | No. CDS | Total Length (Mb) | N50 | %GC | % Complete | CheckM Contam | No. Contig | No. tRNAs | No. 5S rRNA | No. 16S rRNA | No. 23S rRNA |
| --- | --- | --- | --- | --- | --- | --- | --- | --- | --- | --- | --- |
| **Reference MAG** | 1,434 | 1.57 | 145756 | 31.8 | 98.28 | 0 | 5 | 37 | 4 | 4 | 4 |
| **343442** | 1,421 | 1.54 | 75124 | 31.3 | 98.28 | 0 | 35 | 20 | 1 | 1 | 0 |
| **342826** | 1,586 | 1.70 | 122755 | 31.1 | 98.28 | 0 | 29 | 19 | 1 | 1 | 0 |
| **342587** | 1,428 | 1.57 | 103492 | 31.3 | 98.28 | 0 | 27 | 19 | 0 | 1 | 1 |
| **342495** | 1,553 | 1.64 | 90657 | 31.3 | 98.28 | 0 | 31 | 19 | 1 | 1 | 0 |
| **342491** | 1,586 | 1.69 | 95115 | 31.3 | 98.28 | 0 | 30 | 19 | 1 | 1 | 0 |
| **323896** | 1,488 | 1.60 | 110599 | 31.6 | 98.28 | 0 | 35 | 19 | 1 | 1 | 1 |

### Phylogenetic Analysis

A phylogenetic analysis using the full-length 16S rRNA genes revealed BVAB1 belongs in the *Clostridales* family *Lachnospiraceae*, and that *Shuttleworthia satelles* is the closest known relative of BVAB1, though nucleotide identity is only 89-90% (**Figure 1**). Relative to each other, the BVAB1 16S rRNA genes were >99.7% similar and the MAG average nucleotide identity was also high (98.6-99.1%, **Table 3**). At the genome level, the ANI of the BVAB1 MAGs and the *Shuttleworthia satelles* draft genome (GCF000160115) was on average 74.7%. We, therefore, propose a new genus for BVAB1 which represents the phylogenetic placement, the source of these particular MAGs, and the as yet uncultivated nature of this species: *Lachnovaginosum* genomospecies.

**Figure 1.**
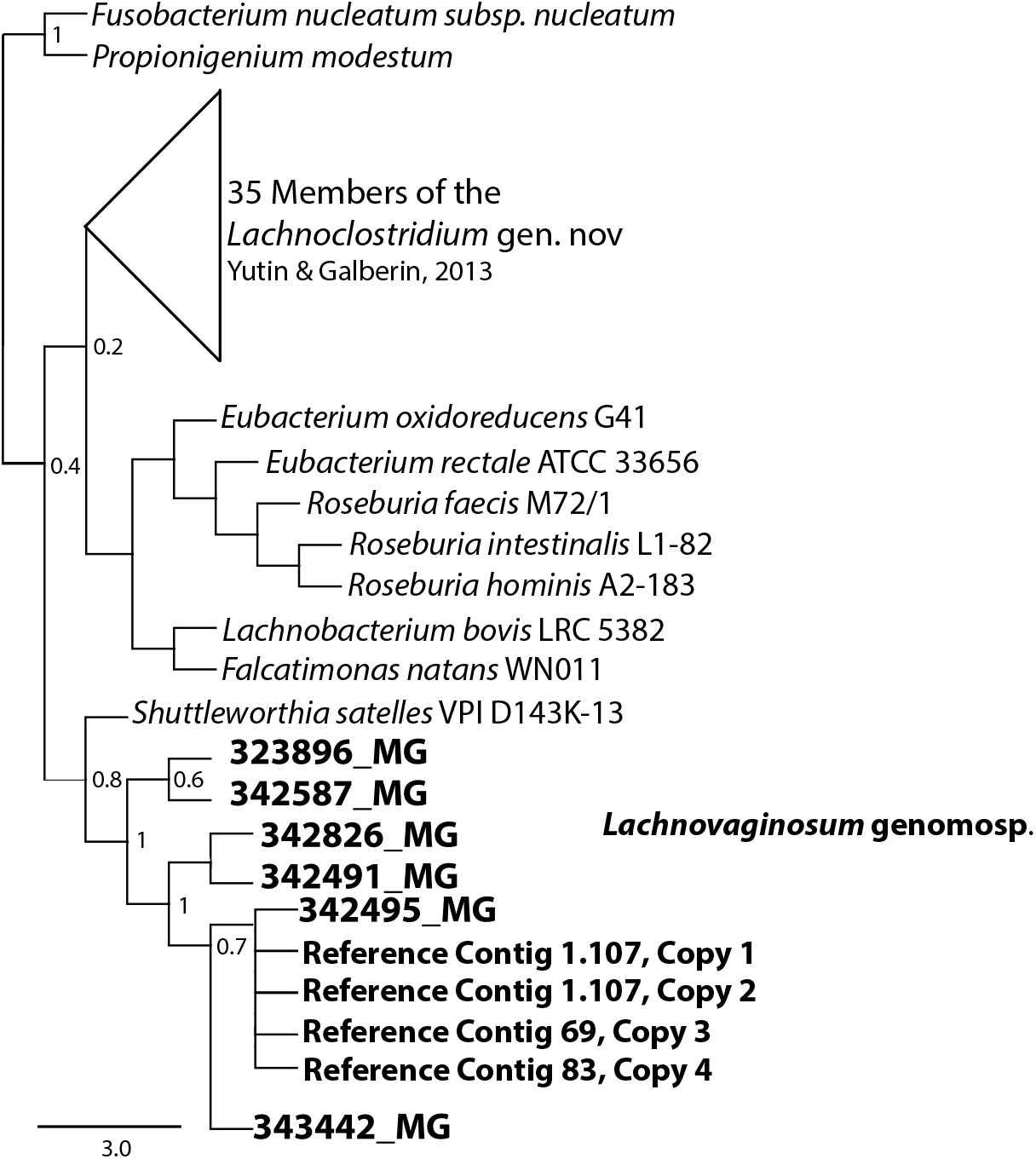
*Lachnovaginosum* genomospecies is related to *Shuttleworthia satelles*. The full-length 16S rRNA gene sequences are 89-90% identical. *Lachnovaginosum* genomospecies sequences, in bold, are >98.6% identical to each other.

**Table 3.** Average Nucleotide Identity (ANI) between each *Lachnovaginosum* genomospecies metagenome-assembled genomes and the *Shuttleworthia satelles* draft genome GCF000160115.

|  | 323896 | 342491 | 342495 | 342587 | 342826 | 343442 | Reference Genome |
| --- | --- | --- | --- | --- | --- | --- | --- |
| **342491** | 99.01 | - | - | - | - | - | - |
| **342495** | 99.11 | 98.88 | - | - | - | - | - |
| **342587** | 99.02 | 98.81 | 98.85 | - | - | - | - |
| **342826** | 98.89 | 99.10 | 99.02 | 99.11 | - | - | - |
| **343442** | 98.88 | 98.93 | 98.92 | 99.09 | 99.01 | - | - |
| **Reference Genome** | 98.83 | 98.60 | 98.86 | 98.86 | 98.73 | 98.80 | - |
| ***Shuttleworthia satelles*** | 74.86 | 74.21 | 74.27 | 74.66 | 75.88 | 74.34 | 74.31 |

### *Specific genomic features of* Lachnovaginosum *genomospecies*

Several genomic features were conserved across the reference genome and the 6 MAGs (**Figures 2** & **3**). Regarding *Lachnovaginosum* genomospecies metabolism, genes encoding transporter for mannose, fructose, and L-ascorbate were identified in all MAGs, as well as a D-lactate dehydrogenase gene but not an L-lactate dehydrogenase gene. This result was unexpected, as the production of D-lactate in the vaginal environment is thought to be a key and somewhat unique feature of certain *Lactobacillus* spp. [42, 43]. Metabolomic data for each sample were explored for the presence of lactate to provide evidence of this potential genomic finding. While lactate was observed in most samples, its abundance was substantially lower than that from representative samples that were dominated by *Lactobacillus crispatus* (CST I), a known D-lactate producer (**Figure 4**). Thus, either the enzyme exhibits lower activity in *Lachnovaginosum* genomospecies or is instead used in the reverse reaction to consume *Lactobacillus* spp.-produced D-lactate to produce pyruvate. This is a strategy described for another vaginal anaerobic Gram-negative coccus, *Veillonella parvula*, which is able to grow on lactate as the sole source of carbon [44]. This would possibly contribute to an increase in vaginal pH, thereby creating a more favorable growth environment. Instead, succinate was the more abundant potential fermentation end product in the communities from which *Lachnovaginosum* genomospecies’ genomes originated. Nonetheless, succinate is associated with bacterial vaginosis, a condition in which *Lachnovaginosum* genomospecies is often found [45]. Interestingly, all *Lachnovaginosum* genomospecies MAGs lack all steps of the tricarboxylic acid cycle. As a consequence, *Lachnovaginosum* genomospecies is certainly unable to produce succinate and it is possible that *Lachnovaginosum* genomospecies produces a substrate another species in the community can convert into succinate. Mannose was also noticeably absent in all of the samples, suggesting that it, too, could be a carbohydrate source for *Lachnovaginosum* genomospecies, which is also supported by the presence of mannose transporter genes on the MAGs (**Figure 3**).

**Figure 2.**
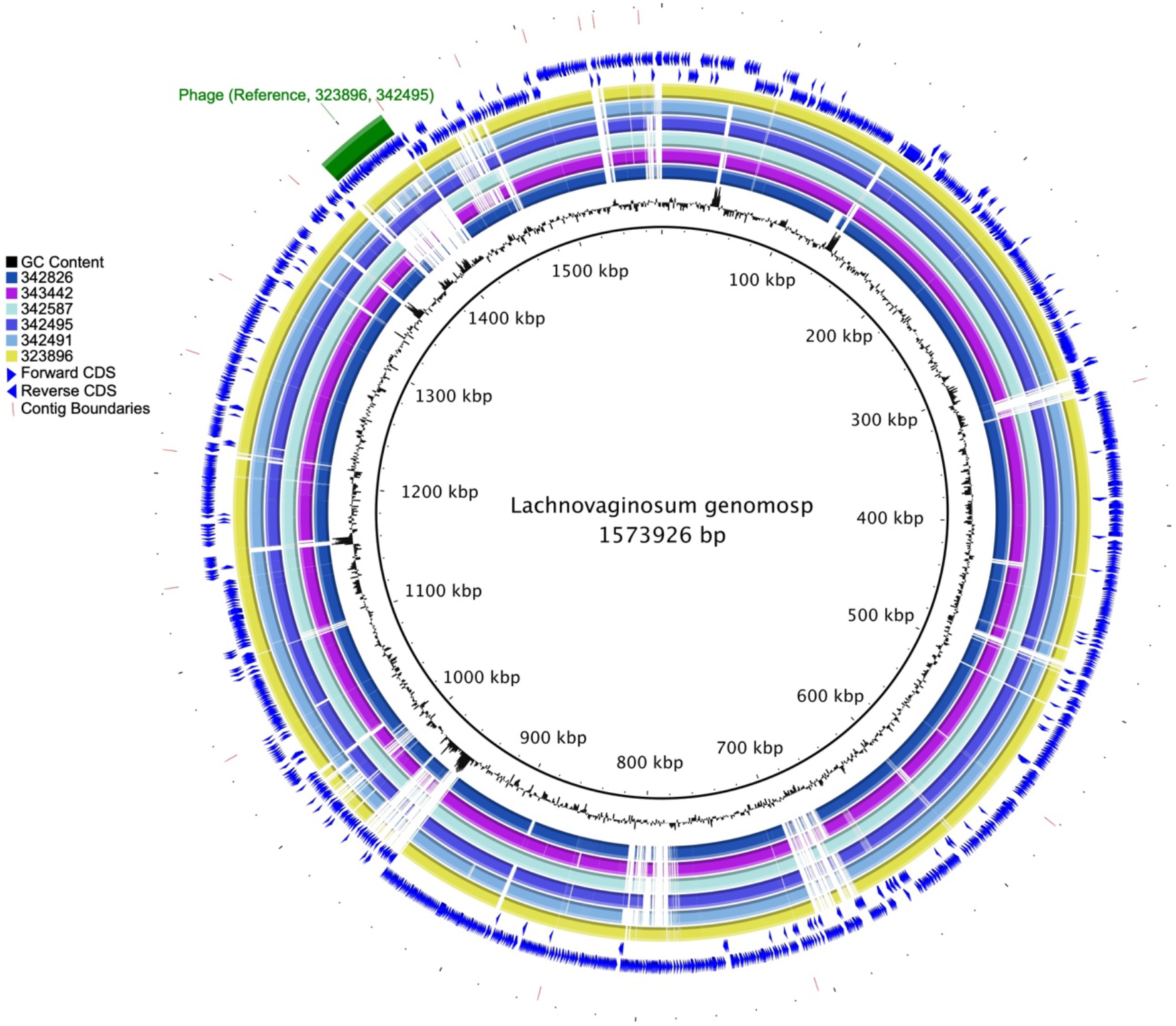
Circle plot comparing the *Lachnovaginosum* genomosp. metagenome-assembled genomes. The MAG sequenced on the Sequel II platform is the reference. Circles are from inside to outside: 1) genome size, 2) GC content, followed by MAGs 3) 342826 4) 343442, 5) 342587, 6) 342495, 7) 342491, 8) 323896 and finally 9) forward-direction coding sequences, 9) reverse-direction coding sequences, 10) phage detected in reference genome (green bar), and 11) contig boundaries for reference genome.

**Figure 3.**
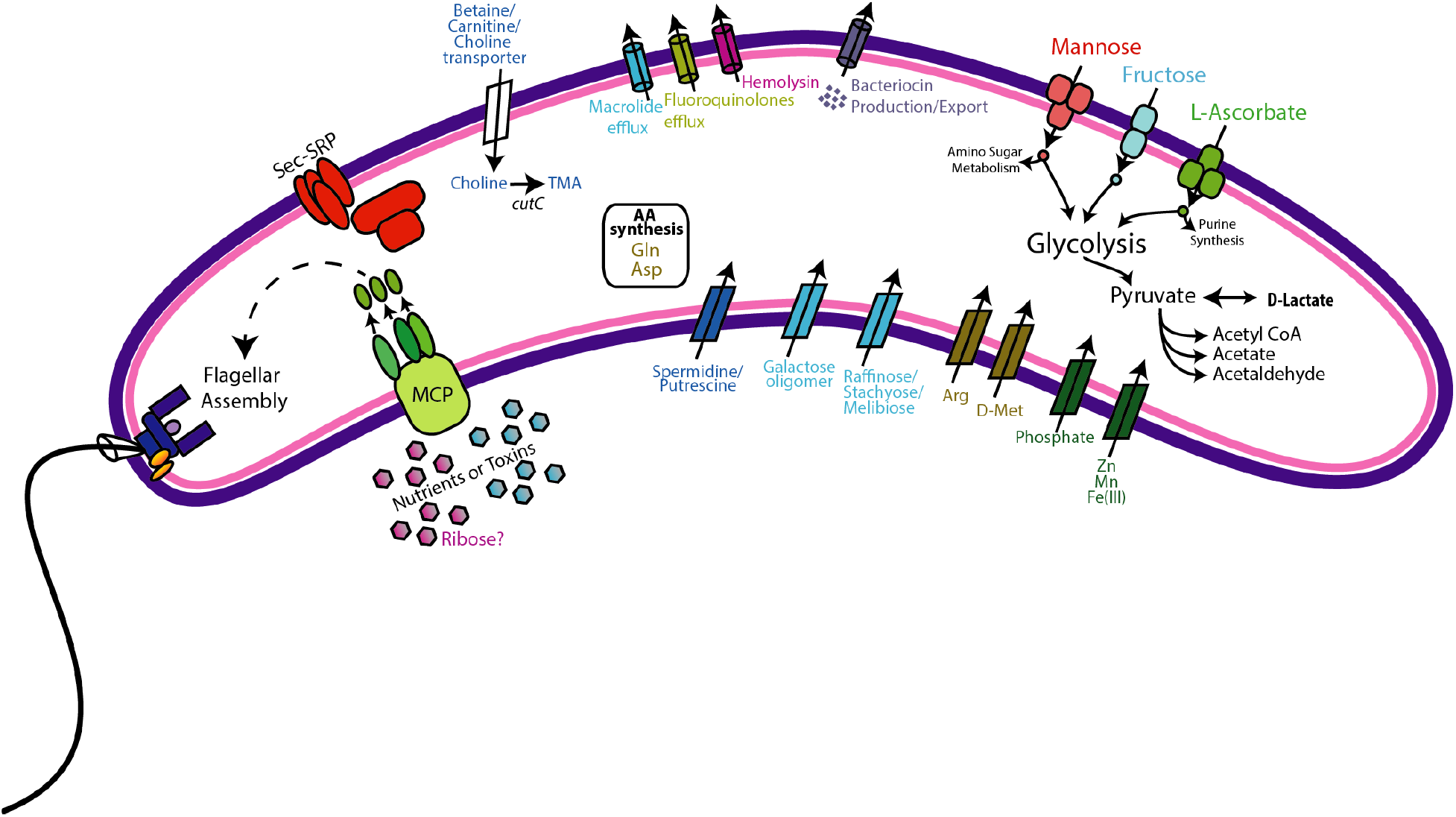
Genomic features of *Lachnovaginosum* genomospecies. Genes coding for the following functions were conserved across all genomes: Methyl-accepting chemotaxis (MCP) and subsequent flagella assembly, Sec-SRP secretion systems, choline import and metabolism, bacteriocin production and export, mannose, fructose, and L-ascorbate transport and metabolism, D-lactate dehydrogenase and biosynthesis of only 2 amino acids.

**Figure 4.**
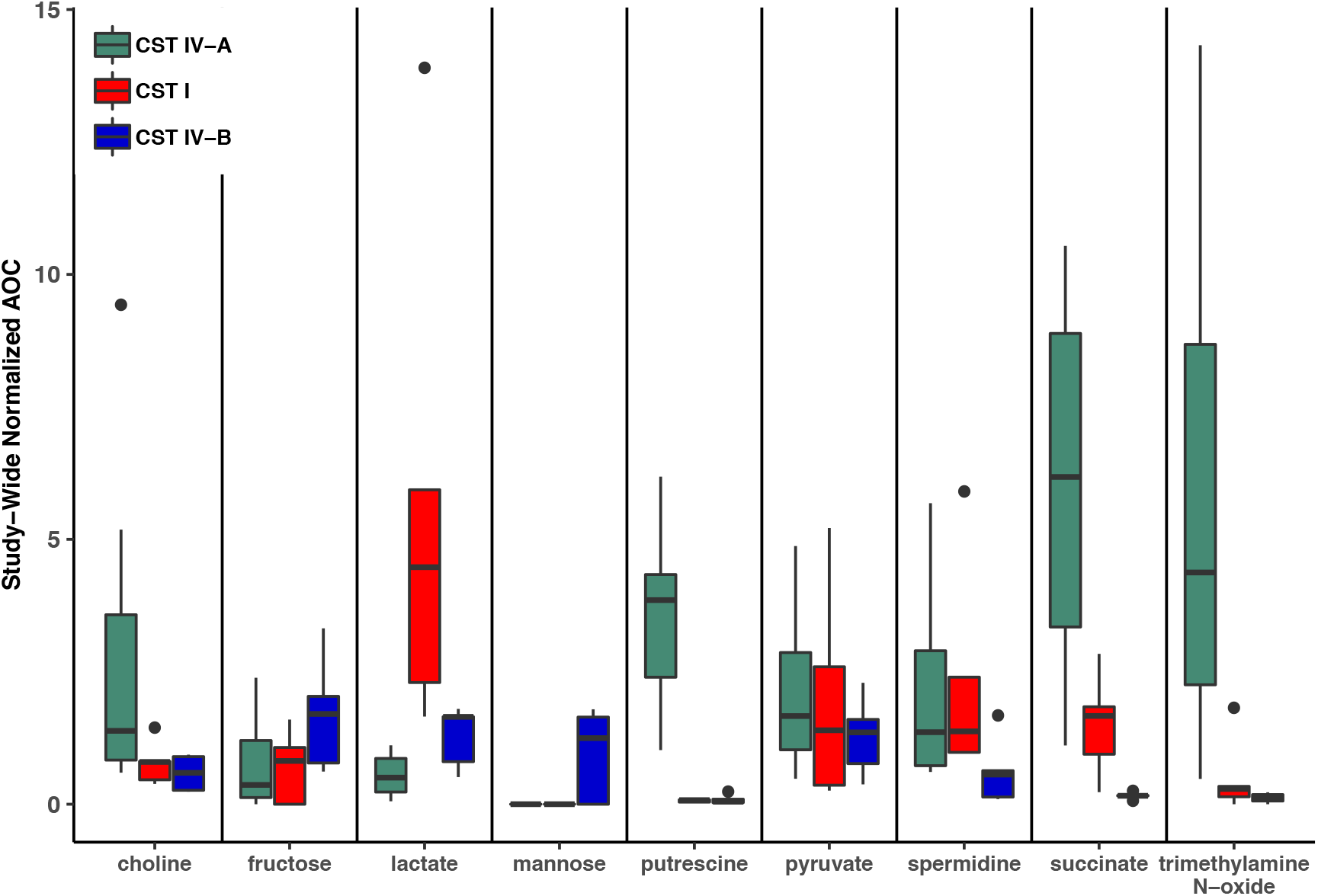
Metabolomic signatures for biochemicals of interest in samples from samples of vaginal community state type (CST) IV-A (relatively high abundances of *Lachnovaginosum* genomospecies), CST I (dominated by *Lactobacillus crispatus*) and CST IV-B (*Gardnerella vaginalis*).

In addition, the full suite of genes required for flagella assembly was observed in all MAGs, as well as genes required for methyl-accepting chemotaxis and the downstream signal process that mediates flagellar response [46]. Flagella have yet to be visually observed on *Lachnovaginosum* genomospecies (**Figure 5**). Other interesting genes include efflux pumps for both macrolides and fluoroquinolones, as well as those encoding hemolysins, which would suggest direct interaction between *Lachnovaginosum* genomospecies and host tissues. Additionally, choline transporters and the *cutC* and *cutD* genes were found in all MAGs, however, the *cutD* gene is absent from the reference MAG. The *cut* genes metabolize choline and produce trimethylamine (TMA) [47], one of the substances believed to be responsible for the fishy odor associated with bacterial vaginosis [48]. TMA was in greater abundance in the metabolomes of these samples relative to those from *Lactobacillus crispatus* or *Gardnerella vaginalis* dominated communities (CST I and CST IV-B) (**Figure 4**).

**Figure 5.**
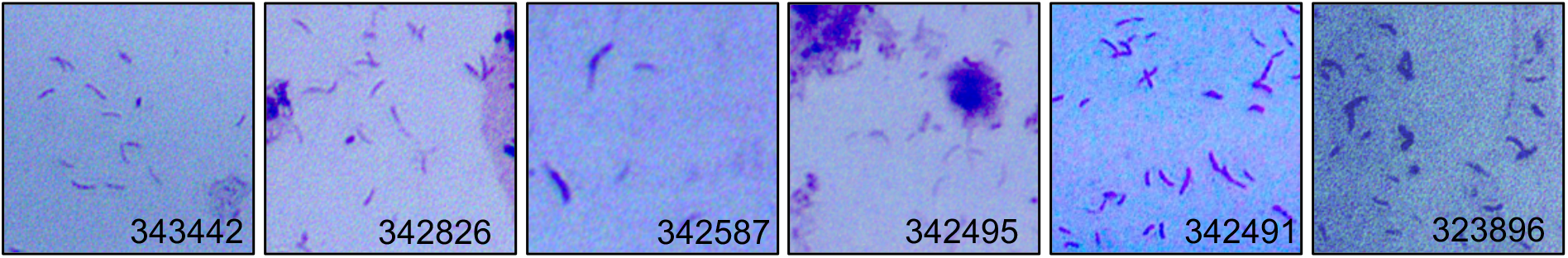
Gram stains of LSVF samples containing >70% *Lachnovaginosum* genomosp. and used in this study. Morphologically *Lachnovaginosum* genomosp. is a curved rod. Images are labeled with sample ID.

Intact phages were detected in 3 MAGs, including the PacBio-produced reference genome (**Table 4**). Best BLAST hits indicate that all detected phages belong to the Siphoviridae family of double-stranded viruses which exhibit both lytic and lysogenic phases. Phages of this family have previously been reported in the vaginal species, *Lactobacillus jensenii* [49].

**Table 4.** Best blast hits and accession numbers of intact phage detected in *Lachnovaginosum* genomospecies metagenome-assembled genomes.

|  | **Intact Phage** | **Accession No.** |
| --- | --- | --- |
| **323896** | Staphy_80 | NC_030652 |
| **342491** | None | - |
| **342495** | Clostr_phiMMP03 | NC_028959 |
| **342587** | None | - |
| **342826** | None | - |
| **343442** | None | - |
| **Reference**<br>**MAG** | Staphy_80 | NC_030652 |

## Conclusions

We present here a collection of *Lachnovaginosum* genomospecies MAGs, previously known as BVAB1, an important member of the human vaginal microbiota associated with bacterial vaginosis and other adverse outcomes. Both the inability to culture this organism as well as its detection via a small region of the 16S rRNA gene sequence have led a limited understanding of its contributions to the vaginal microbiome and implications for its effect on reproductive health. We have shown the potential of this microbe to be motile in the environment most likely in response to nutritional cues, to resist antibiotics using efflux systems, and to contribute to the fishy odor characteristic of bacterial vaginosis. Further work on the characterization of *Lachnovaginosum* genomospecies will be necessary to establish its role in the vaginal microbiome.

## Declarations

### Availability of data and materials

The datasets and supplementary materials generated and/or analyzed in the current study are available in the FigShare repository under doi: 10.6084/m9.figshare.8194595.

## Acknowledgments

The authors are grateful to the University of Maryland School of Medicine Genome Resource Center and Elizabeth O’Hanlon for providing the Gram-stained images.

